# Local field potential sharp waves with diversified impact on cortical neuronal encoding of haptic input

**DOI:** 10.1101/2023.11.10.566542

**Authors:** Sofie S. Kristensen, Henrik Jörntell

## Abstract

Cortical sensory processing is greatly impacted by internally generated activity. But controlling for that activity is difficult since the thalamocortical network is a high-dimensional system with rapid state changes. Therefore, to unwind the cortical computational architecture there is a need for physiological ‘landmarks’ that can be used as frames of reference for computational state. Here we use a waveshape transform method to identify conspicuous local field potential sharp waves (LFP-SPWs) in the somatosensory cortex (S1). LFP-SPW events triggered short-lasting but massive neuronal activation in all recorded neurons with a subset of neurons initiating their activation up to 20 ms before the LFP-SPW onset. In contrast, LFP-SPWs differentially impacted the neuronal spike responses to ensuing tactile inputs, depressing the tactile responses in some neurons and enhancing them in others. When LFP-SPWs coactivated with more distant ECoG-SPWs, suggesting an involvement of these SPWs in global cortical signaling, the impact of the LFP-SPW on the neuronal tactile response could change substantially, including inverting its impact to the opposite. These cortical SPWs had similar overall activity patterns as reported for hippocampal SPWs and may be a biomarker for a particular type of state change that possibly involves both hippocampus and neocortex.

## Introduction

How neocortical information processing is configured remains open to various alternative interpretations. In recent years, multiple observations suggest that representations of information are widespread across neuron populations [1-3] and that individual neurons within certain constraints carry complementary information [4-6], even when they are located literally right next to each other and are provided with the exact same input [5, 7]. However, also the time-continuous variations in internal brain state impact how the individual neurons represent the information. The exact same tactile input pattern can trigger a wide variety of responses in individual primary somatosensory (S1) cortical neurons [8]. The response variations are due to internal state variations in the neocortical neuronal network globally, which can be demonstrated by weakly perturbing either a remote area of the cortex [9, 10] or the output of the hippocampus [11]. These response variations in individual neurons can be explained as a consequence of subtle changes in the state of the neocortex globally, which in turn can alter the number of “open” network pathways supplying the recorded neuron with tactile information. Hence, when interpreting the response of a neuron to any given input, it is important to also control for the current internal cortical state when the input is provided.

But controlling for that internal activity is difficult since the thalamocortical network is high-dimensional, perpetually active and its state can change at a high pace. There is therefore a value in identifying potential markers for such internally generated state changes. Here we apply a template-based signal extraction method to identify a wide category of events that in many respects resembles the hippocampal sharp wave, but which is recorded as spontaneous local field potential sharp waves (LFP-SPWs) at about mid-depth in S1 neocortex.

The hippocampal sharp wave (Hipp-SPW) [12] is a prominent field potential signal in the dendritic field layer of the hippocampal pyramidal neurons, stratum radiatum. When recorded in the pyramidal cell layer, these Hipp-SPWs typically have higher-frequency ripples superimposed. The ripples can be relatively easily detected after band-pass filtering and have been associated with many high-level brain functions, such as replay and recall [13-16]. The Hipp-SPW has been linked to activity changes in the neocortex in different ways [17-20]. For example, Hipp-SPWs can trigger similar EEG-SPWs as demonstrated for the prefrontal cortical region [20]. Interestingly, a recent paper reported that ‘contrary to the model in which SWRs arise ‘spontaneously’ in the hippocampus, neocortical activation often precedes SWRs and may thus constitute a trigger event in which neocortical information seeds associative reactivation of hippocampal ‘indices’’ [21].

We found our S1 cortex LFP-SPWs to have similar overall activation characteristics as reported for Hipp-SPWs. LFP-SPWs also strongly correlated with similar sharp waves recorded at the cortical surface using ECoG electrodes (‘ECoG-SPWs’). As the ECoG-SPWs were recorded some distance away from the LFP-SPWs, this suggests that these SPWs represent some type of non-local cortical signal. In the spontaneous activity, both types of SPWs triggered a massive drive on the neuronal spiking activity. We found that preceding LFP-SPWs greatly impacted the S1 neuronal tactile responses. However, the impact was differentiated, in some neurons they caused a reduction whereas in other neurons they caused a substantial enhancement of the evoked response. The ECoG SPWs, which occurred more rarely than the LFP-SPWs (but always coincided with them), could in turn substantially modify the impact that the LFP-SPW had on the tactile response.

## Methods

### Surgical procedure

Adult Sprague-Dawley rats (N=16, male sex, weight 306-420) were maintained under general anesthesia through a continuous infusion of ketamine and xylazine (ratio 20:1) mixed with Ringer acetate and glucose. The anesthetic mixture was administered as a continuous infusion through an intravenous catheter in the right femoral vein (approximately ∼5 mg/kg per hour). The catheter was inserted by making an incision in the inguinal area of the hindlimb. Before inserting the catheter, animals were sedated with isoflurane (3% mixed with air for 60–120 s) and given an intraperitoneal injection of ketamine/xylazine (ratio 15:1) to induce anesthesia. While anesthetized, a small part of the skull (4×4 mm) was removed to expose the primary somatosensory (S1) cortex. The craniectomy extended from the coordinates 1 mm rostral to −3 mm caudally and 2-4 mm laterally relative to bregma (**Figure 1A**). An ECoG-electrode was placed on the surface of the cortex at the caudal end of the craniectomy. To prevent dehydration of the brain, the exposed brain area was covered in a thin layer of agarose (0.03 g/ml dissolved in physiological saline). Apart from recording SPWs, the ECoG signal was used to monitor the level of anesthesia by characterizing occurrences sleep spindles mixed with epochs of more desynchronized activity, an indication of sleep [22]. Note that ketamine/xylazine anesthesia has previously been shown to not affect the order of neuronal recruitment of a sheet of layer V neurons in spontaneous brain activity fluctuations and evoked responses as compared to the awake condition, suggesting that the neocortical network otherwise may work close to normal [23]. Adequate anesthesia was further ensured by noxious pinching of the hind paw to confirm absence of withdrawal reflexes. Animals were sacrificed with pentobarbital (140 mg/kg IV) by the end of the experiment. All procedures related to animal experiments were approved by the Local Animal Ethics Committee of Lund, Sweden in advance (permit M13193-2017).

**Figure 1.**
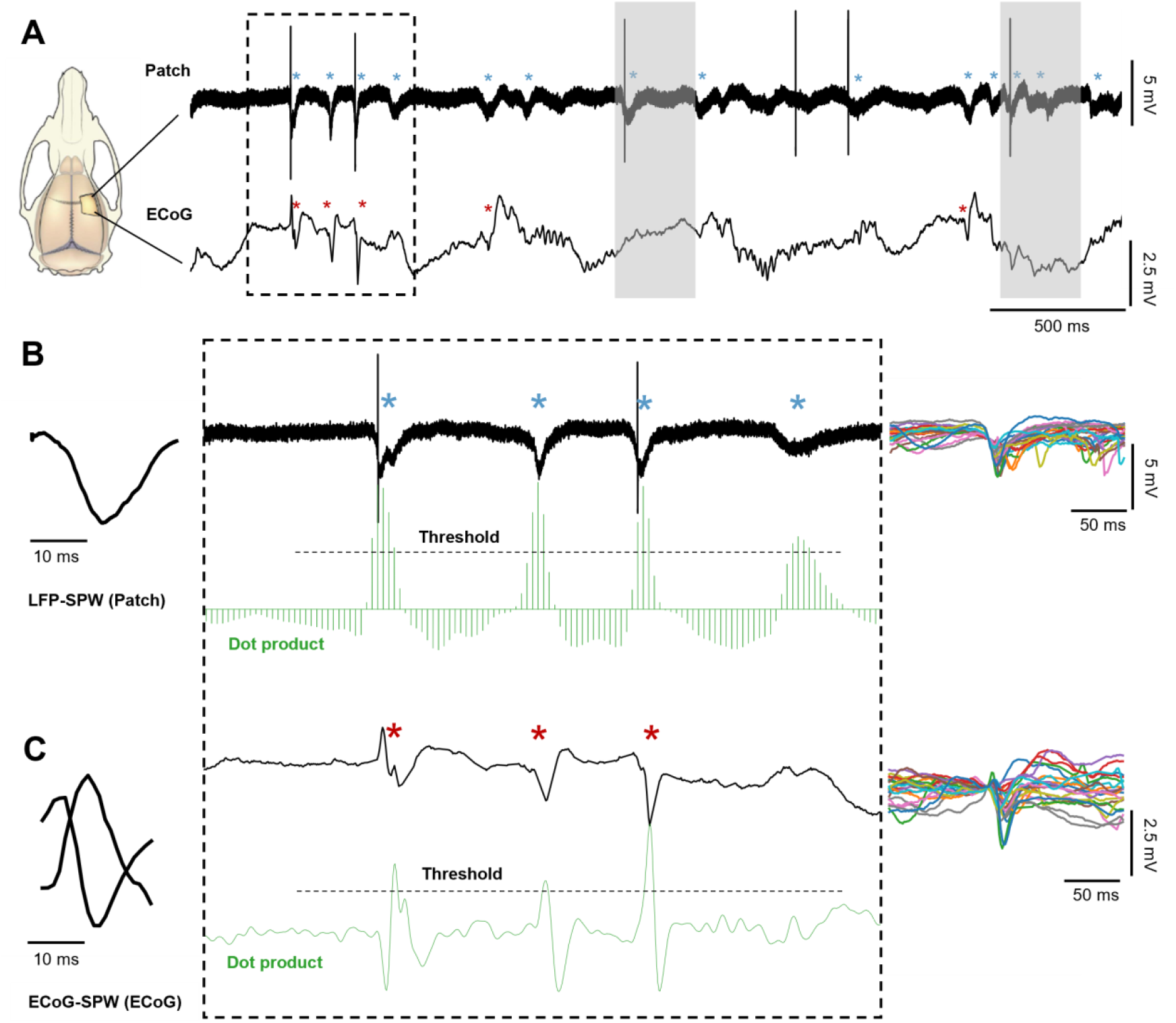
Properties of LFP and ECoG sharp waves. (A) Left: LFP and ECoG recording sites inside and outside the forelimb S1 region, respectively. Right: Raw recording data from a sample experiment. The asterisks indicate LFP sharp waves (LFP-SPWs) and ECoG sharp waves (ECoG-SPWs), respectively. Grey shading indicates periods of skin stimulation on the second digit. (B) Left: LFP-SPW template. Middle: Zoom-in on a segment of the raw traces in A. Below, in green, are the equivalent traces of the dot product, obtained by multiplying the templates with the raw recordings across each time step. The SPW-detection threshold is indicated for each dot product trace as black dashes lines. Right: 25 superimposed LFP-SPWs to illustrate their variability in appearance. (C) Same as B, but for the ECoG recording.

### Recordings

Electrocorticographic recordings (ECoG) were made from the surface of the cortex at the rostral end of the craniectomy, approximately 4 mm laterally and 3 mm caudally to the bregma using a silver ball electrode (Ø 250 um at the cortex contact surface; no filters were applied, amplifier cut-off 50 kHz). Extracellular recordings of individual neurons were made with patch clamp pipettes in the loose-patch current clamp recording mode from the forepaw region of the S1, at the coordinates −1.0 – 1.0 mm relative to bregma. The location of the S1 was also estimated by evoked LFPs in response to stimulation of the second digit of the forepaw. Borosilicate glass capillaries were used to pull the patch pipettes to 10–30 MΩ using a Sutter Instruments (Novato, CA, USA) P-97 horizontal puller. Pipettes were filled with an electrolyte solution composited of (in mM) potassium-gluconate (135), HEPES (10), KCl (6.0), Mg-ATP (2), EGTA (10) titrated to7.35–7.40 pH using 1 M KOH. The pipette was inserted into the cortex with an electrical stepping motor at a rate of 0.002 mm/s and neurons were recorded between cortical layers III and V. Unfiltered patch electrode recording data was digitized at 100 KHz using CED 1401 mk2 hardware and Spike2 software (CambridgeElectronic Design, CED, Cambridge, UK). The ECoG recording was digitized at 1 kHz using the same system.

### Stimulation

We wanted to analyze the effect of LFP-SPWs on the evoked spike responses to haptic stimulation. We have previously shown that most S1 neurons have unique responses to electrotactile spatiotemporal activation patterns mimicking mechanical skin-object interactions [2, 3, 5].The patterns consisted of 5-33 pulses with intensities of 0.5 mA and durations of 0.14 ms (DS3 Isolated Stimulator, Digitimer, UK) delivered through four pairs of needle electrodes. Needle pairs were inserted into the volar side of the second digit of the forepaw with 2-3 mm distance between each pair. Patterns were delivered in pre-defined random order and lasted between 200 ms and 340 ms and were separated by about 1.8 s (random intervals). Each pattern was repeated 50 times in one experiment. These were thus the exact same set of spatiotemporal activation patterns we have used in earlier publications, but since we in the current paper only cared about the earliest part of the response evoked by the first stimulation pulses for each pattern, we simply considered pooled responses from all stimulation patterns and refer the interested reader to the original papers in order to view the full patterns.

### Responsiveness to stimulation

Before pooling the responses, we calculated the responsiveness to each of the eight stimulation patterns for individual neurons. We constructed peristimulus time histograms (PSTHs) of the spike responses across all 100 repetitions of a stimulation pattern (in a few cases only 50 repetitions). The spike responses in the 200 ms time window preceding the onset of the stimulation were binned at 2 ms time bins and used to calculate the mean spike frequency and standard deviation (SD) of the baseline activity. If the average poststimulus activity (30 ms) exceeded the baseline activity by more than two SDs, then it was counted as being responsive. Stimulation patterns that did not evoke a response according to this definition, were excluded from the pooled responses of the neuron used for further analysis. Only neurons with responses to at least 3/8 of the patterns (i.e. at least 150 repetitions of tactile inputs) were included in the analysis.

### Extracting neuron spikes, LFP-SPWs and ECoG-SPWs

Neuron spike events were extracted and time-stamped using an in-house software employing a template based spike identification [24]. Local field potential sharp waves (LFP-SPW) and ECoG sharp waves (ECoG-SPW) were identified using a waveshape transform approach, which is related to continuous wavelet transform but employs a user-defined waveshape rather than a wavelet. An LFP-SPW waveshape was based on visually identified LFP-SPWs, which had a latency to peak of 10-15 ms and a duration of 20-30 ms. The LFP waveshape vector was multiplied with the vector of the time continuous patch recording data, with a stride of 500 data sample points (5 ms), across the entire recording (typically 30-60 mins). At each stride, we used the dot product of the transform to define if an SPW was present by setting a threshold. We manually set the threshold of the dot product for each neuron recording individually, so that as many visually identified events as possible could be detected without including other, less structured variations in baseline activity. We removed detected events if they were detected less than 10 ms apart to avoid the same event being detected twice. The same method was used to detect ECoG-SPWs in the ECoG recording, except that here we used two waveshape templates; one with a positive voltage deflection and one with a negative voltage deflection. Here, the waveshape was multiplied with a stride of one data sample point (1 ms).

### Spike responses to tactile stimulations, LFP-SPWs or ECoG-SPWs

To visualize the relationships between the spike activity and the tactile stimulations, the LFP-SPWs and the ECoG-SPWs, we constructed Kernel Density Estimations (KDEs) of the spike responses. Each triggered trace consisted of a time series of spikes. Each individual spike response, occurring at a specific time point relative to the trigger (which was a tactile stimulation or a SPW), was convolved with a 2 ms Gaussian kernel. Then all the convolved responses of that category were summed to obtain a KDE of the response around the trigger point.

## STATISTICAL ANALYSIS

### Classifying the spike responses triggered by spontaneous SPWs

For the spike responses to the SPWs, the intensity and latency time of the spike activity that preceded the onset of the LFP-SPWs were used to define three distinct groups; neurons with no preceding activity, neurons with short preceding activity, and neurons with longer preceding activity. To sort neurons into these groups, we calculated the normalized baseline KDE activity and its SD in a time window starting 100 ms to 20 ms in the pre LFP-SPW interval separately for each neuron. The time window was chosen because the earliest preceding activity was observed to precede the LFP-SPW onset by 20 ms. If a neuron did not have any spike activity exceeding its baseline by two SDs in the 20 ms time window preceding LFP-SPW onset, the neuron would fall into the category “no preceding activity”. If a neuron exceeded baseline activity by two SDs within 7 ms preceding LFP-SPW onset, it would be categorized as “short preceding activity”. If a neuron exceeded baseline activity by two SDs at more than 7 ms preceding LFP-SPW onset, it would be categorized as “longer preceding activity”. The same procedure was repeated using ECoG-SPWs as the trigger.

To statistically analyze if ECoG-SPWs evoked more spikes than LFP-SPWs, the amplitudes of KDEs were measured for each individual neuron in each group and compared with Welch’s t-test. Before running the test, the distributions of the two groups were checked for normality by plotting histograms. *Levene*’s test for homogeneity of variance failed which is why Welch’s t-test was chosen instead of a paired student’s t-test.

### LFP-SPWs impact on spike responses evoked by tactile stimulation

To analyze the effect of LFP-SPWs on the evoked spike responses for individual neurons, all recording traces were divided into two groups for each neuron individually. The first group of recording traces were the ones in which stimulations were not preceded by an LFP-SPW in the 100 ms time window preceding stimulation onset. The second group consisted of traces where stimulations were preceded by an LFP-SPW in this time window. To compare the evoked responses to the stimulations for these two groups of traces, the number of spikes occurring in the 30 ms post stimulation time window was counted for each trace in each group. This resulted in two distributions of ordinal data. The Mann-Whitney U test was used to test for a statistical difference between the two groups of traces (traces preceded by and traces not preceded by an LFP-SPW) in terms of the number of spikes in each trace. If the test result was significant, and the difference between the normalized sum of the evoked spikes was negative, we classified the neuron as having a response that was “depressed” by the SPW. If the difference between the normalized sum of the evoked spikes instead was positive, we classified the neuron as being “excited” by the SPW. If the test result was not significant, we classified the neuron as having “no effect” from the preceding SPW. Effect size was calculated with: 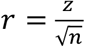

### ECoG-SPWs effect on the LFP-SPWs impact on evoked responses

The same method as described above was used to compare evoked responses preceded by LFP-SPWs in the 100 ms pre stimulus time window but with or without a coinciding ECoG-SPW. Also here, neurons were classified as “depressed”, “excited”, or with “no effect” depending on the significance of the test. In this case, depressed would indicate that the coinciding ECoG-SPW reduced the evoked response, and vice versa for excitation. Only neurons that had 15 or more traces per category were included in the corresponding analysis.

### Wavelet analysis

A wavelet analysis was performed on three different groups of SPWs for one example experiment. A continuous wavelet transform (Complex morlet wavelet, base cycle=‘your parameter’, between 1–350 Hz) was computed and the wavelets were normalized to the integral of the modulus of the wavelet function in time domain. This was done for all LFP-SPWs, all LFP-SPWs coinciding with ECoG-SPWs and all ECoG-SPWs separately. The averages of each of the three analyses were obtained and plotted as frequency spectrograms. The raw ECoG signal was upscaled to the same sampling frequency as the raw patch signal before performing the wavelet analysis.

## Results

Our aim was to investigate the effect of LFP-SPWs and ECoG-SPWs, which both stood out as salient signals in our recordings (**Figure 1A**), on tactile-evoked neuronal spiking responses. LFP-SPWs were recorded with the same patch electrode that was also used to record unitary neuronal spikes, and the ECoG signal was obtained from the surface of the cortex at least 2 mm away from the patch electrode. Neuron recordings were made across layers III-V (depth 0.432-1.094 mm, except one neuron at depth 0.330 mm), which corresponded to the depths where the LFP-SPWs were found to be the most prominent. A total number of 65 neurons were recorded from 16 rats with neuronal responses evoked by haptic stimulation delivered to the second digit of the forepaw. 48 neurons passed the inclusion criterion for analysis (Methods).

### Shape and frequency analysis of SPWs

**Figure 1A** illustrates a four-second-long segment of a parallel patch and ECoG recording from one example neuron. Light blue asterisks indicate LFP-SPWs in the patch recording and dark red asterisks indicate ECoG-SPWs in the ECoG recording. Left part of **Figure 1B and 1C** shows the wavelet templates used to detect the presence of LFP-SPWs and ECoG-SPWs, in the patch recording and the ECoG recording, respectively. Middle part of **Figure 1B and 1C** shows a zoom in on 0.5 seconds of the segment together with the dot product traces that resulted from multiplying the waveshape templates with the raw data. Right part of **Figure 1B and 1C** illustrates that both LFP-SPWs and ECoG-SPWs were highly variable in their appearance (latency-to-peak and peak amplitude for example, see **Table 1**), which naturally made it impossible to capture all the occurrences of either type of event. According to our detection method (Methods), LFP-SPWs occurred more than 2 times more frequently than ECoG-SPWs (**Table 1**). ECoG-SPWs occurred almost exclusively in coincidence with LFP-SPWs, which is illustrated in **Figure 2B** where the occurrence of LFP-SPWs was triggered on ECoG-SPWs. **Figure 2A and 2C** illustrate the inter-event-intervals (IEIs) for LFP-SPWs (light blue) and ECoG-SPWs (dark red) respectively, from the same example neuron recording. IEIs of SPWs were not dissimilar from inter-spike-intervals (ISIs) (**Figure 2G**) in shape, i.e. ISIs and IEIs both had a tendency towards a sharp peak followed by a log-normal style of distribution, as previously described for a wide set of neurons [25, 26], but the SPWs occurred at a much slower overall rate. What can also be observed in **Figure 1A** is that spikes sometimes coincided with the LFP-SPWs (**Table 1**), but spikes could also occur outside these events.

**Table 1:** Frequencies of occurrence of LFP-SPWs and ECoG-SPWs across all 48 neurons. Coincidence frequency between LFP-SPWs and ECoG SPWs, and between LFP-SPWs and spikes. Quantification of the variety of LFP-SPW configurations.

|  |  |
| --- | --- |
| **LFP-SPW** | 3.3 Hz (1.8 Hz) |
| **ECoG-SPW** | 1.4 Hz (0.8 Hz) |
| **LFP-SPW with ECoG-SPW** | 33.8 % (16.7 %) |
| **LFP-SPW with spike** | 36.9 % (20.2 %) |
| **LFP-SPW amplitude** | -1.3 mV (0.7 mV) |
| **LFP-SPW latency to peak** | 11.6 ms (5 ms) |

**Figure 2.**
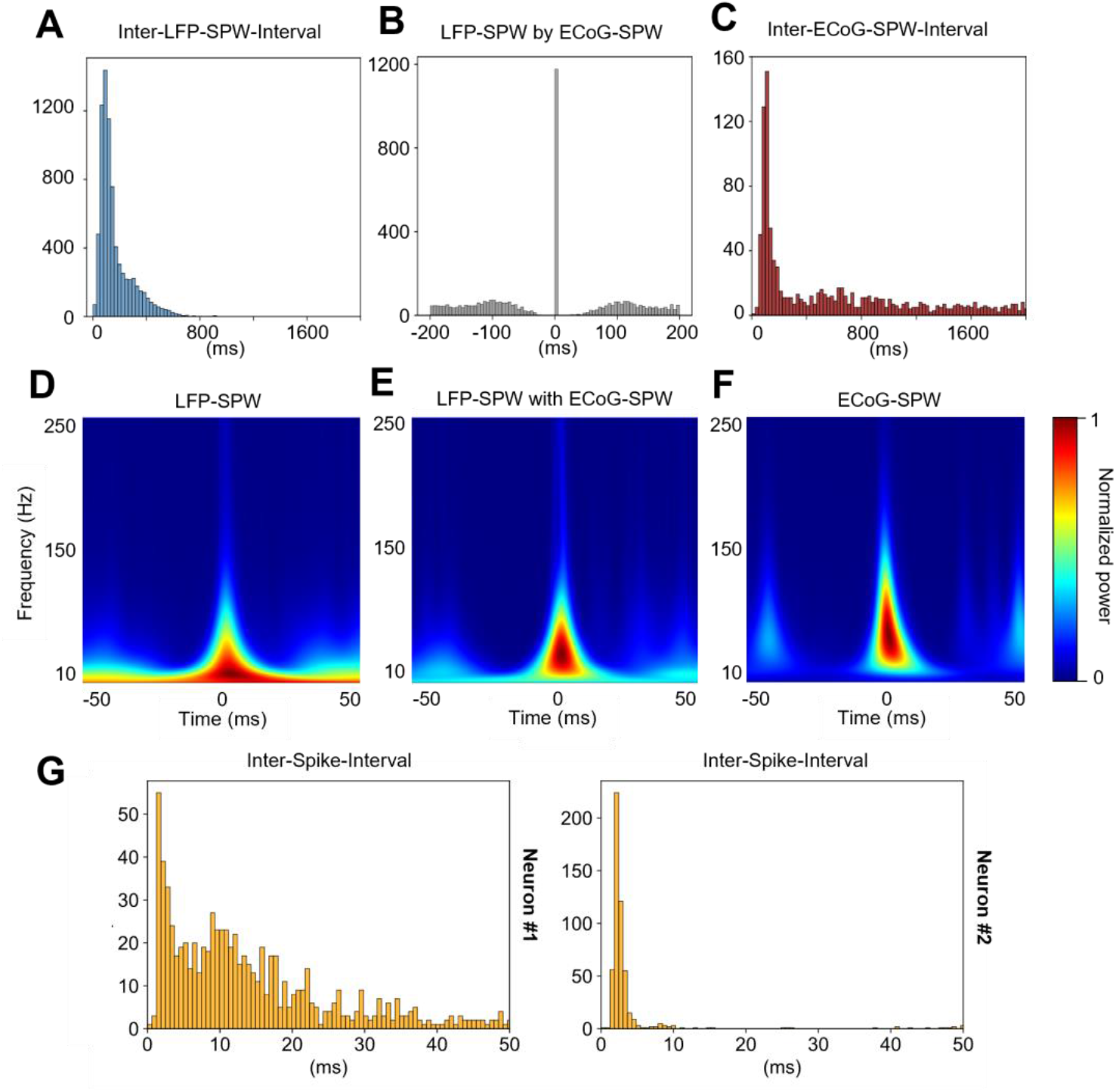
Frequency analysis of LFP-SPWs, ECoG-SPWs and spikes. (A) Inter-event intervals of LFP-SPWs. (B) Occurrences of LFP-SPWs triggered by occurrences of ECoG-SPWs. (C) Inter-event intervals of ECoG-SPWs. (D) Wavelet spectogram of all LFP-SPWs in one experiment. (E) Wavelet spectogram of LFP-SPWs coinciding with ECoG-SPWs in the same experiment. (F) Wavelet spectogram of ECoG-SPWs in the same experiment. (G) Interspike intervals (ISIs) for two example neurons, for comparison with the Inter-event intervals in A and C. Note the much shorter time scales of the ISI histograms.

### The effect of LFP-SPWs on evoked spike responses

The first part of the analysis was centered in our main question, namely, if LFP-SPWs had an impact on the tactile-evoked spike responses. To analyze this, we divided all tactile-evoked recording traces of a neuron into two classes, depending on whether they were preceded or not preceded by an LFP-SPW in the 100 ms prestimulus time window. **Figure 3A** illustrates example raw traces and KDEs of all responses for a sample neuron where the presence of a preceding LFP-SPW (in the grey box time window) resulted in a depression of the tactile-evoked response (measured in the red time zone). **Figure 3B** instead illustrates a neuron where the preceding LFP-SPW resulted in an excitation of the tactile-evoked response.

**Figure 3.**
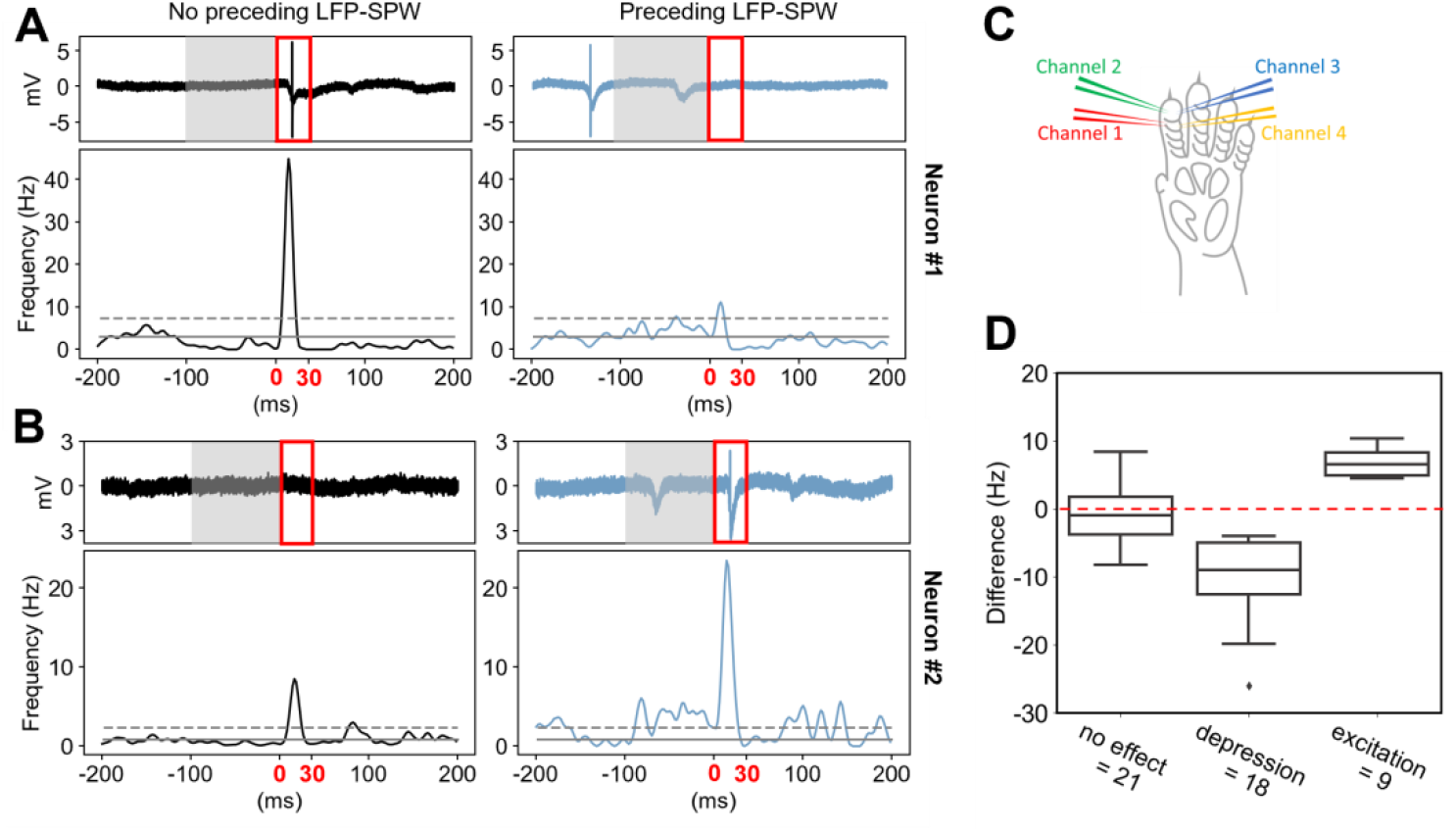
Differential impact of LFP-SPW on tactile evoked responses across neurons. (A) A neuron for which the response to tactile stimulation was depressed when it was preceded by an LFP-SPW. The response is shown as a sample raw trace and as a KDE plot of the average evoked spike response. Left panel corresponds to responses that were not preceded by an LFP-SPW (in the time window shaded in grey). Right panel corresponds to the evoked spike responses when the tactile stimulation was preceded by a LFP-SPW. Red box highlights the time window where the evoked spike responses were counted. (B) Similar display as in A, but for a neuron in which the spike response was enhanced when there was a preceding LFP-SPWs (compare left with right panel). (C) Electrotactile stimulation sites. (D) Summary of LFP-SPW impacts on the tactile evoked spike responses. Neurons were either found to not have statistically significant effect of preceding LFP-SPWs, to have a significant depression, or to have a significant spike response enhancement.

To statistically analyze the differences between the two classes of responses for every neuron individually, we used Mann-Whitney U as a test. The test showed a significant difference in the evoked responses between the populations of traces with a preceding LFP-SPW and the population of traces without a preceding LFP-SPW for 30 out 48 neurons (see supplementary data for details from individual tests). Of these 30 neurons, 21 showed a decrease in evoked spikes when preceded by an LFP-SPW, indicating that the presence of LFP-SPWs within 100 ms pre stimulus onset were depressing the evoked spike response. Nine of the neurons with a significant difference, showed an increase in evoked spikes when preceded by an LFP-SPW, indicating excitation of evoked responses. In 17 of the 21 significantly affected neurons, the effect size of the difference was small (*r* < 2), in three neurons the effect was small-moderate (*r* > 2, < 3), and in one neuron the effect was moderate (*r* > 3) (exact effect sizes are reported in the Supplementary Data). The difference in spike frequency in the 30 ms response windows is displayed as box plots in **Figure 2D** separately for the three groups of neurons (no significant effect, significant depression, and significant excitation).

### The impact of SPWs on neuronal spiking

SPWs had a major impact on neuronal spiking (**Figure 4A-C**). When we used SPWs in the spontaneous activity to trigger the spike occurrences, the corresponding PSTHs (here shown as KDEs) illustrated massive spike responses (**Figure 4B,C**). Interestingly, a subset of our neurons had a build-up of activity starting before the occurrence of the SPWs. This suggests they contributed to eliciting the SPWs (as has been suggested for hippocampal neurons for Hipp-SPWs [27]). We could separate the neurons into three categories (**Figure 4B**), neurons without preceding spiking activity (N=13), neurons that had spiking activity that preceded the LFP-SPW for a short time window (7 ms or less; N=22) and neurons that that had spiking activity in a long time window (more than 7 ms, maximally 20 ms was observed) preceding the onset of the LFP-SPW (N=13). For the ECoG SPW, the same categories were observed but individual neurons did not necessarily fall into the same category as for the LFP-SPW. When using ECoG-SPWs as the trigger, a total number of 14 neurons showed no preceding activity, 25 neurons showed short preceding activity and 9 neurons showed long preceding activity (Figure 3C). The ECoG-SPWs in general evoked a more powerful spike response than the LFP-SPWs. In 34 neurons, the peak amplitude for the ECoG-SPW KDEs was higher compared to the amplitude for the LFP-SPW KDEs (**Figure 4D)**. Welch’s t-test indicated that the mean amplitude in the ECoG-SPW group (62.4 ± 34.9 Hz) was significantly higher than the mean amplitude in the LFP-SPW group (41.7 ± 29.5 Hz) with a difference of 20.6 (95% CI; 15.2 to 26.1); t(47) = 3.1, *p* <0.01 one tailed. Overall, the intense neuron responses indicated that the local network was in a different state when the SPWs occurred.

**Figure 4.**
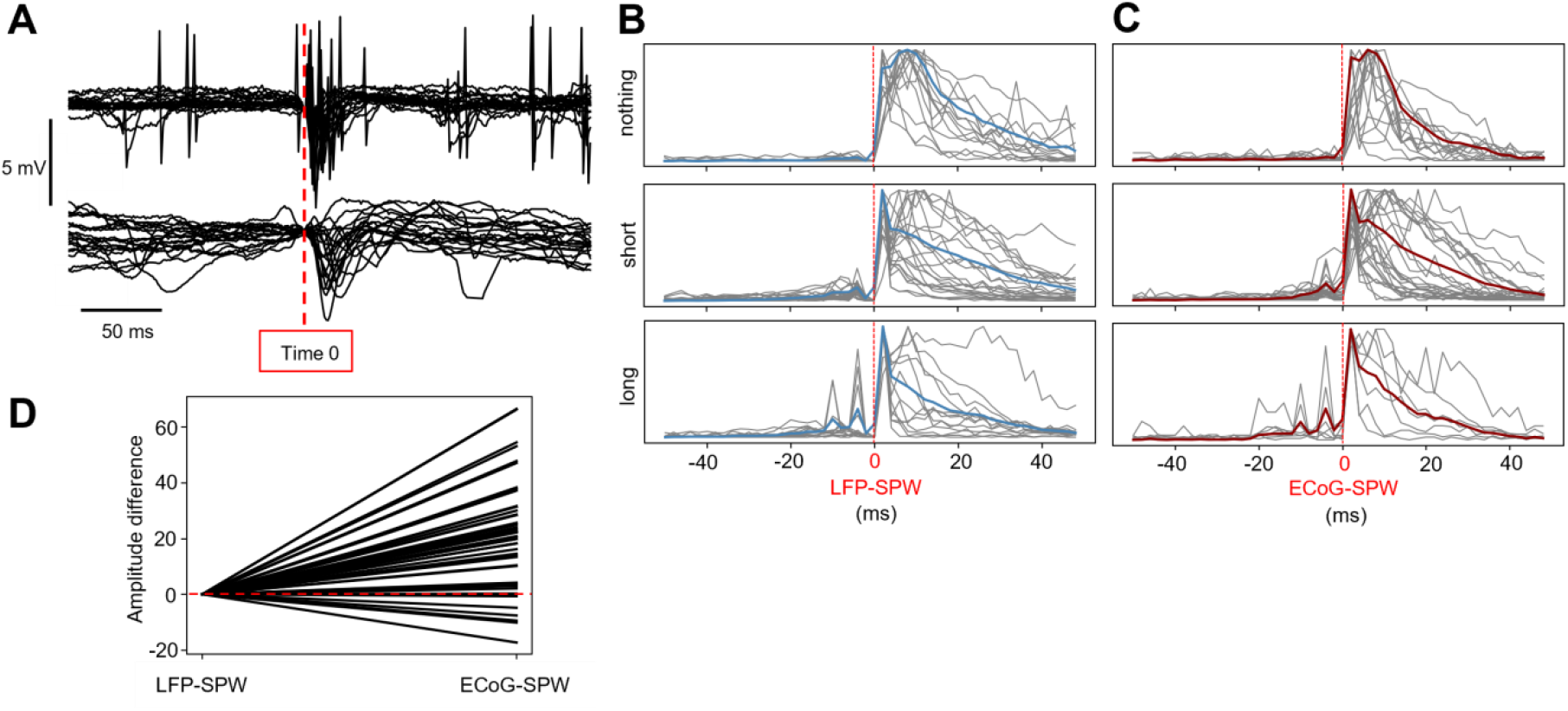
Spike responses triggered by LFP-SPWs and ECoG-SPWs (A) Superimposed raw data traces for a sample neuron, to illustrate spike responses around LFP-SPWs (top) and concomitant ECoG-SPWs (bottom). (B) Normalized SPW-triggered spike responses sorted by neurons with an absence of a preceding spike response, a short preceding response (up to 7 ms) and a long preceding response (up to 20 ms before the onset of the SPW). Individual neuron responses are shown as KDEs in grey, the average KDEs are shown in light blue (LFP-SPWs). (C) Similar display as in A but with ECoG-SPWs as the trigger, the average KDE curves are shown in dark red. (D) The difference in peak amplitudes between the LFP-SPW triggered spike responses and the ECoG triggered spike responses across all individual neurons.

### ECoG-SPWs affected the LFP-SPWs impacts on the evoked spike responses

Considering the more powerful effect of ECoG-SPWs on spontaneous spiking, we next analyzed if the ECoG-SPWs affected the impact that LFP-SPWs had on evoked responses when they coincided. We divided all recording traces for each individual neuron into two classes; traces where preceding LFP-SPWs coincided with an ECoG-SPW and traces where preceding LFP-SPWs did not coincide with an ECoG-SPW. Example KDEs for normalized traces from the two groups are shown in the second and thirds columns of **Figure 5A-C** for three example neurons as dark blue traces (no coinciding ECoG-SPW) and dark red traces (with a coinciding ECoG-SPW). The first column with black traces illustrates traces that were not preceded by any LFP-SPW or ECoG-SPW.

**Figure 5.**
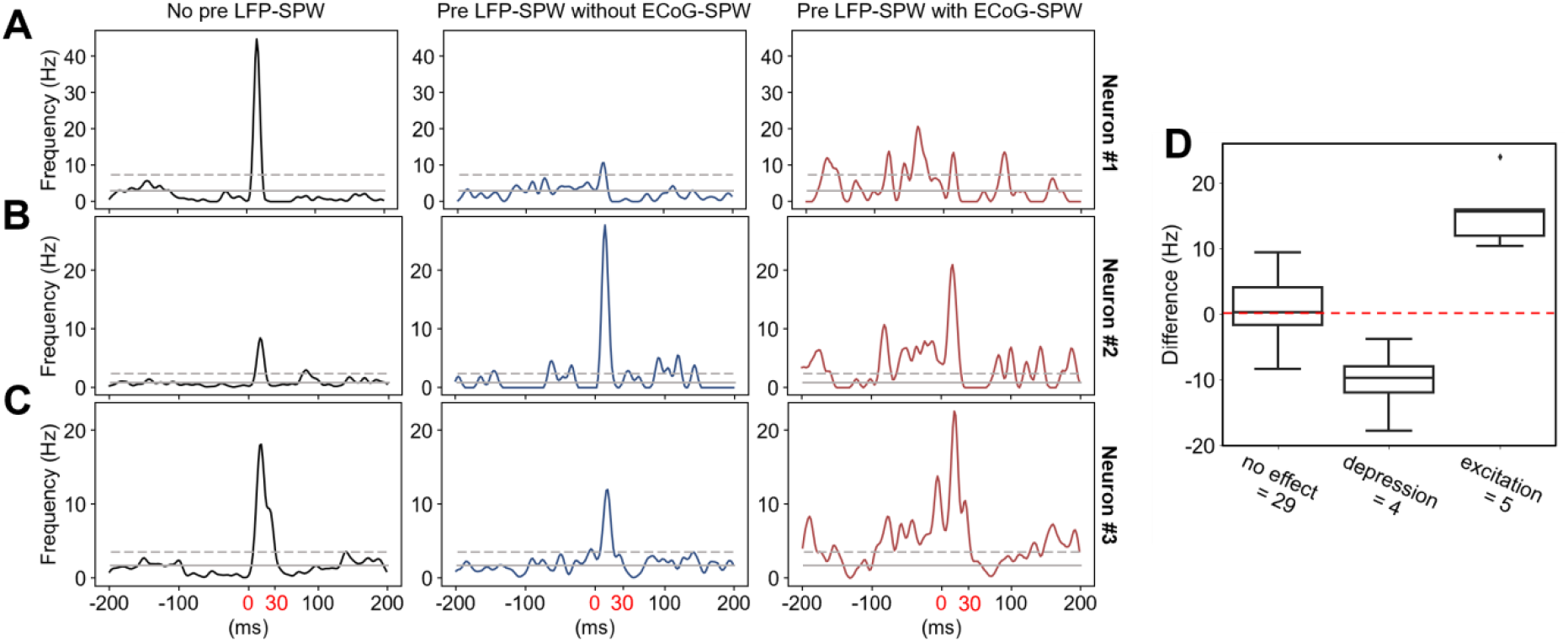
Effect of coinciding ECoG-SPWs on the LFP-SPW-mediated impacts on evoked spike responses. (A) An example neuron with a substantial depressing impact of the preceding LFP-SPWs on the evoked response (dark blue KDE trace) but in which the coincidence of an ECoG-SPW with the LFP-SPW did not affect that impact (dark red KDE trace), despite a marked enhancement in the prestimulus spiking activity. (B) A neuron where the preceding LFP-SPW enhanced the response (dark blue trace) but where the coincident ECoG-SPW+LFP-SPW depressed that impact (dark red trace). (C) A neuron where the preceding LFP-SPW depressed the response but where the coincident ECoG-SPW+LFP-SPW inverted that impact to instead enhance the response above its control level (black trace to the left). (D) Effect of coinciding ECoG-SPWs on the LFP-SPW-mediated impacts on the evoked spike responses (i.e. differences between dark blue KDEs and dark red KDEs across all neurons).

As before, we compared the evoked responses in the traces from the two classes using the Mann-Whitney U statistical test (10 neurons with too few coinciding traces were excluded from the analysis). In this case, it indicated a significant difference for nine of the neurons. The effect size of the difference for these nine neurons showed five neurons with a small effect (*r* < 2) and four neurons with a small-moderate effect (*r* > 2, < 3) (see Supplementary Material for more detailed test results). Of the nine neurons that showed a significant difference in the evoked responses, four neurons showed a reduction by the coincidence of an ECoG-SPW (depressed) and five neurons showed an enhancement (excited) as shown in **Figure 5D**.

### Control analysis

Since ECoG-SPWs elicited more intense spike response than LFP-SPWs (**Figure 4**), and affected the impact of the LFP-SPWs on the evoked responses (**Figure 5**) the next question we asked was whether the mere presence of a spike in the LFP-SPW had a similar boosting effect as the ECoG-SPW. Again we divided all recording traces for each individual neuron into two classes, LFP-SPWs with spikes and LFP-SPWs without any coinciding spike (i.e. no spike occurring between the onset of the SPW and two times the latency to peak) and compared the groups using Mann-Whitney U. Example KDEs for the two classes of traces of one neuron is shown in **Figure 6A**. Note the increased spike activity in the 100 ms time window preceding the stimulation, indicated by the grey box. As before, we analyzed the evoked spikes in the 30 ms post-stimulation onset time window (indicated by red numbers in **Figure 6A**). In this analysis, only 2/48 neurons had a significant difference between the evoked responses in the two groups (**Figure 6B**). The effect size for both neurons was small (*r* < 3). See Supplementary Material for details on Mann-Whitney U tests and effect sizes.

**Figure 6.**
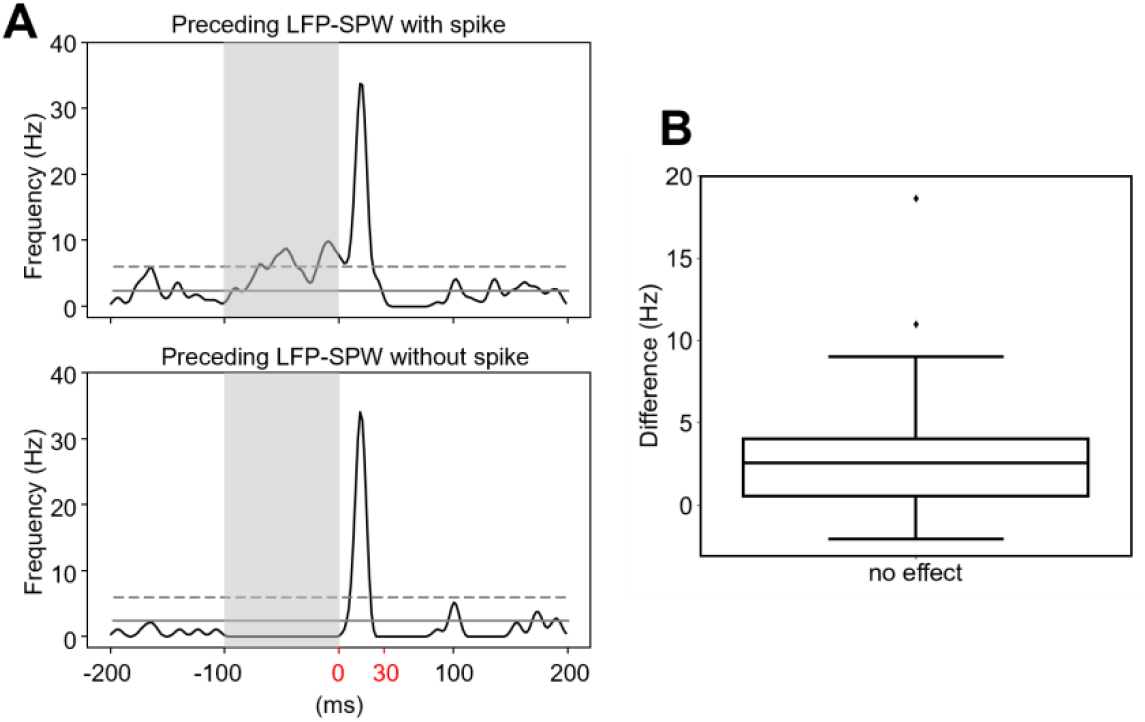
Control analysis if the presence of spiking affected the LFP-SPWs impacts on the evoked responses. (A) KDEs of the spike responses when the LFP-SPW contained spike responses (top) or not (bottom). (B). Quantitative analysis indicated that the presence of spike responses in the LFP-SPWs did not significantly impact the evoked response except for the two outliers above.

## Discussion

Here we reported on a particular type of neocortical LFP event, the LFP SPW, which we found to have a relatively large and diversified impact on the neuronal responses evoked by tactile stimulation. This finding could be explained by that the LFP-SPW was associated with a significant state change, at least in the local circuitry that mediated the tactile input to the recorded neuron. The LFP-SPW often coincided with an ECoG-SPW recorded some distance away, which suggests that at least a subset of the LFP-SPWs was associated with a more widespread cortical state change. The activation dynamics of our cortical SPWs were similar to SPWs previously recorded in the hippocampus, and previous studies showed that hippocampal SPWs trigger SPW-like EEG-signatures at least in the prefrontal cortex [20] or possibly the other way around [21]. Hence, the LFP-SPWs may potentially be part of a widespread coordination of cortical neuron activity.

We found that the LFP-SPW could have opposing impacts on the tactile-evoked responses in different neurons. This is in line with previous observations that even adjacent neurons can encode given sensory inputs in a diversified manner [5, 7], or have complementary patterns of spontaneous activity [4, 24, 28]. In relation to the LFP-spike relationship more generally, a previous study concluded that there is a ‘…substantial heterogeneity in the timing and strength of spike-EEG relationships and that these relationships became more diverse during visual stimulation compared with the spontaneous state.’ [29]. Our results for the LFP-SPWs are hence in line with that more general EEG relationship, and further emphasize the diversity of the relationships between individual neurons and population level features in the brain activity.

The state change signaled by the LFP SPWs appeared to be quite different when the LFP-SPW coincided with an ECoG-SPW. This was indicated by that the impact that the LFP-SPW had on the tactile evoked response in a neuron (depression or excitation) could even be inverted if the LFP-SPW coincided with an ECoG-SPW (**Figure 5**). Although spontaneous SPWs triggered massive neuronal spike responses on average (**Figure 4**), the mere presence or absence of spiking in the preceding LFP-SPW overall did not alter the impact it had on the evoked response (**Figure 6**). This indicates that the spikes themselves were not good indicators of a state change, in contrast to the SPWs.

The neuronal spiking activity was powerfully driven by the SPWs. This suggests that the SPWs were associated with powerful synaptic drive to the local neurons. Many neurons also had activity profiles that indicated that they were activated well in advance of the SPW initiation (**Figure 4**). This may in addition suggest that the LFP-SPWs were a result of a rapid but gradual build-up of activity in recurrent excitatory loops that at least partly involved the local neurons – similar to the current interpretations of how Hipp–SPWs arise [13, 27]. Since Hipp-SPWs are associated with concomitant intense activity in the neocortex, including the S1 [18], this in turn raises the question if it is the hippocampus that paces the neocortex, is it the other way around [21], or is it predominantly a distributed contribution across both the hippocampus and the neocortex?

### Similarities with Hipp-SPWs

In many respects the LFP-SPWs displayed similarities with Hipp-SPWs. The time-frequency signatures are very similar (Figure 2; cf. Buzsáki [13], Liu, et al. [30]; Petersen, et al. [31]), though the lack of superimposed ripples made the cortical LFP-SPWs occupy a lower frequency range than the Hipp-SPWs. Although the overall occurrences of Hipp-SPWs are often not analyzed in detail in the literature, many comparisons can be made based on raw data illustrations. The first similarity is their approximate duration [13, 20]. A second similarity is their rate of occurrence – Hipp-SPWs have been reported to occur at more than 1 Hz [14] and example illustrations indicate rates in the order of up to 2 Hz [20]. This is less than the 3.3 Hz we found for LFP-SPWs, but in line with the rate of our ECoG-SPWs. However, as discussed below, our detection method using the waveshape transform may be more sensitive and thus missing fewer SPW events. In addition, Hipp-SPWs have been reported to be less frequent in awake active state than in awake relaxed state, sleep and anesthesia [14] as was the case in our experiments. Also, the cortical SPWs could occur in bouts of 2-4 SPWs in a series with much shorter intervals between them (see **Figure 1**). Unfortunately, there is not a lot of analysis on SPW IEIs *in vivo* in the literature, but raw data illustrations (i.e. Figure 1 in Buzsáki [13] for example) suggest this is the case also for hippocampal SPWs.

### What do the SPWs represent and how common are they?

Why were the rates of occurrence of the LFP-SPWs lower than for the ECoG-SPWs? This might in fact possibly have been due to event detection issues – SPWs were easier to detect in LFPs, because of the more precise nature of the signal from the patch than from the silver ball electrode, and because ECoG-SPWs could often be initially positive and then negative, which made them harder to detect. But this highlights another problem that has not been addressed in the literature – the detection threshold for SPWs will vary not only depending on the analysis method but also on what recording method is used. This is likely to apply to hippocampal recordings as well – one method to detect Hipp-SPWs that includes high-speed ripples has been to band-pass filter the field potential data to above 50 Hz and then set the detection threshold to 5 SDs [27]. It is possible that such an approach will be at risk of missing SPWs that do not have a significant part of their signal power above that frequency threshold. This in turn comes back to the question whether SPWs have a strictly unitary character or if these types of events instead are much more gradable. Recent data suggests that Hipp-SPWs are highly variable and gradable (where the reported ripple-free SPWs from stratum radiatum in fact have characteristics that are very similar to the cortical LFP-SPWs, **Figure 2**) [32]. Our detection method did not rely on band-pass-filtering but instead used the dot product obtained with template waveshapes. Even though also our method used an arbitrary detection threshold for the SPWs (**Figure 1**) their highly variable amplitudes and temporal signatures suggest that the SPWs are extensively gradable, perhaps all the way down to being in continuum with overall baseline activity. Then it may be the case that SPWs occur at a much higher rate than reported in the literature – for example, their lower incidence reported in active animals than in quiet awake animals, as well as sleep [14], could possibly be because they are less well synchronized in the active awake state. Lower synchrony could in turn be due to that there is more active processing in the cortex [33] and hippocampus [34]. Alternatively, it may be that the higher baseline activity may tend to drown SPW-like phenomena and thus making it more difficult to detect them.

### Concluding remarks

Future studies should begin to address the underlying circuitry mechanisms that drive these SPWs in greater detail. Only then can we begin to understand what they represent more specifically. So far perhaps the strongest indicator is that it has been found that Hipp-SPWs correlate with familiarity in visual scenes [15] or recollection of visual scenes [35]. But given that there are also neocortical SPW events, could these also represent some type of local familiarity signal? And the fact that we observe them also under anesthesia, both in S1 in the present paper and in the hippocampus [27], where information integration is unlikely to be coherent over longer periods of seconds or more, could they represent some type of time-local information-processing completion? The path towards addressing such questions lies in a better understanding of their circuitry-level mechanistic underpinnings and what they correspond to for the cortical information processing mechanisms in general.

## Data and code availability

The recording data and analysis code will be made freely available on the Figshare server on publication of the paper.

## Acknowledgments

We thank Kaan Kesgin for inventing the dot product approach to identify the LFP sharp waves. We thank Dr. Jonas M.D. Enander for allowing us to use some of his data from the publication doi: 10.1016/j.celrep.2019.02.099. This research was funded by the EU Horizon 2020 research and innovation program under grant agreement No 861166 (INTUITIVE).

