## Supplementary material for "Local field potential sharp waves with diversified impact on cortical neuronal encoding of haptic input"

### 1.A Mann-Whitney U test results 1/3 - No LFP-SPW vs. LFP-SPW (all)

```
[1] "neuron:  AFNR0002"

   Wilcoxon rank sum test with continuity correction

data:  var by group
W = 6970.5, p-value = 0.4171
alternative hypothesis: true location shift is less than 0

[1] "neuron:  AFNR0003"

   Wilcoxon rank sum test with continuity correction

data:  var by group
W = 8429, p-value = 1.412e-11
alternative hypothesis: true location shift is less than 0

[1] "neuron:  AGNR0002"

   Wilcoxon rank sum test with continuity correction

data:  var by group
W = 8345.5, p-value = 0.003011
alternative hypothesis: true location shift is less than 0

[1] "neuron:  AGNR0003"

   Wilcoxon rank sum test with continuity correction

data:  var by group
W = 21851, p-value = 3.661e-06
alternative hypothesis: true location shift is less than 0

[1] "neuron:  AGNR0004"

   Wilcoxon rank sum test with continuity correction

data:  var by group
W = 26914, p-value = 0.05839
alternative hypothesis: true location shift is less than 0

[1] "neuron:  AGNR0005"

   Wilcoxon rank sum test with continuity correction

data:  var by group
W = 37692, p-value = 0.004502
alternative hypothesis: true location shift is greater than 0

[1] "neuron:  AKNR0000"
```

```
   Wilcoxon rank sum test with continuity correction

data:  var by group
W = 22440, p-value = 0.006678
alternative hypothesis: true location shift is less than 0

[1] "neuron:  AKNR0001"

   Wilcoxon rank sum test with continuity correction

data:  var by group
W = 20584, p-value = 0.03128
alternative hypothesis: true location shift is less than 0

[1] "neuron:  AKNR0004"

   Wilcoxon rank sum test with continuity correction

data:  var by group
W = 1343.5, p-value = 0.006988
alternative hypothesis: true location shift is greater than 0

[1] "neuron:  AKNR0005"

   Wilcoxon rank sum test with continuity correction

data:  var by group
W = 15464, p-value = 0.01918
alternative hypothesis: true location shift is less than 0

[1] "neuron:  ALNR0003"

   Wilcoxon rank sum test with continuity correction

data:  var by group
W = 24234, p-value = 0.006411
alternative hypothesis: true location shift is less than 0

[1] "neuron:  ALNR0004"

   Wilcoxon rank sum test with continuity correction

data:  var by group
W = 10679, p-value = 3.565e-05
alternative hypothesis: true location shift is less than 0

[1] "neuron:  ALNR0006"

   Wilcoxon rank sum test with continuity correction

data:  var by group
W = 43534, p-value = 0.0005313
alternative hypothesis: true location shift is less than 0

[1] "neuron:  ANNR0003"

   Wilcoxon rank sum test with continuity correction

data:  var by group
W = 10428, p-value = 0.1451
alternative hypothesis: true location shift is less than 0

[1] "neuron:  ANNR0005"

   Wilcoxon rank sum test with continuity correction
```

```
data:  var by group
W = 11205, p-value = 0.2191
alternative hypothesis: true location shift is less than 0

[1] "neuron:  JANR0001"

	Wilcoxon rank sum test with continuity correction

data:  var by group
W = 9422.5, p-value = 0.1748
alternative hypothesis: true location shift is greater than 0

[1] "neuron:  JANR0002"

	Wilcoxon rank sum test with continuity correction

data:  var by group
W = 3189, p-value = 0.04187
alternative hypothesis: true location shift is greater than 0

[1] "neuron:  JANR0004"

	Wilcoxon rank sum test with continuity correction

data:  var by group
W = 36214, p-value = 0.5867
alternative hypothesis: true location shift is less than 0

[1] "neuron:  JANR0005"

	Wilcoxon rank sum test with continuity correction

data:  var by group
W = 3257.5, p-value = 0.08632
alternative hypothesis: true location shift is less than 0

[1] "neuron:  JANR0006"

	Wilcoxon rank sum test with continuity correction

data:  var by group
W = 15936, p-value = 0.003672
alternative hypothesis: true location shift is less than 0

[1] "neuron:  JANR0007"

	Wilcoxon rank sum test with continuity correction

data:  var by group
W = 14312, p-value = 6.705e-05
alternative hypothesis: true location shift is less than 0

[1] "neuron:  JANR0008"

	Wilcoxon rank sum test with continuity correction

data:  var by group
W = 15496, p-value = 0.00418
alternative hypothesis: true location shift is less than 0

[1] "neuron:  JANR0009"

	Wilcoxon rank sum test with continuity correction

data:  var by group
```

```
W = 14079, p-value = 0.01075
alternative hypothesis: true location shift is less than 0

[1] "neuron:  JANR0010"

   Wilcoxon rank sum test with continuity correction

data:  var by group
W = 16953, p-value = 0.06321
alternative hypothesis: true location shift is less than 0

[1] "neuron:  JANR0011"

   Wilcoxon rank sum test with continuity correction

data:  var by group
W = 12894, p-value = 2.096e-09
alternative hypothesis: true location shift is less than 0

[1] "neuron:  JANR0012"

   Wilcoxon rank sum test with continuity correction

data:  var by group
W = 13739, p-value = 5.059e-06
alternative hypothesis: true location shift is less than 0

[1] "neuron:  JANR0017"

   Wilcoxon rank sum test with continuity correction

data:  var by group
W = 13981, p-value = 0.0003216
alternative hypothesis: true location shift is less than 0

[1] "neuron:  JBNR0003"

   Wilcoxon rank sum test with continuity correction

data:  var by group
W = 3052, p-value = 0.1477
alternative hypothesis: true location shift is less than 0

[1] "neuron:  JBNR0005"

   Wilcoxon rank sum test with continuity correction

data:  var by group
W = 4513, p-value = 0.06749
alternative hypothesis: true location shift is less than 0

[1] "neuron:  JDNR0001"

   Wilcoxon rank sum test with continuity correction

data:  var by group
W = 15583, p-value = 0.1723
alternative hypothesis: true location shift is greater than 0

[1] "neuron:  JDNR0002"

   Wilcoxon rank sum test with continuity correction

data:  var by group
W = 16372, p-value = 0.005657
```

```
alternative hypothesis: true location shift is greater than 0

[1] "neuron:  JENR0001"

   Wilcoxon rank sum test with continuity correction

data:  var by group
W = 6736.5, p-value = 0.5262
alternative hypothesis: true location shift is greater than 0

[1] "neuron:  JENR0002"

   Wilcoxon rank sum test with continuity correction

data:  var by group
W = 1208, p-value = 0.009548
alternative hypothesis: true location shift is greater than 0

[1] "neuron:  JENR0003"

   Wilcoxon rank sum test with continuity correction

data:  var by group
W = 13288, p-value = 0.1512
alternative hypothesis: true location shift is greater than 0

[1] "neuron:  JENR0004"

   Wilcoxon rank sum test with continuity correction

data:  var by group
W = 4450.5, p-value = 0.2433
alternative hypothesis: true location shift is less than 0

[1] "neuron:  JGNR0006"

   Wilcoxon rank sum test with continuity correction

data:  var by group
W = 18741, p-value = 0.02887
alternative hypothesis: true location shift is greater than 0

[1] "neuron:  JGNR0007"

   Wilcoxon rank sum test with continuity correction

data:  var by group
W = 13902, p-value = 0.009957
alternative hypothesis: true location shift is less than 0

[1] "neuron:  JGNR0008"

   Wilcoxon rank sum test with continuity correction

data:  var by group
W = 14435, p-value = 0.2867
alternative hypothesis: true location shift is less than 0

[1] "neuron:  JGNR0009"

   Wilcoxon rank sum test with continuity correction

data:  var by group
W = 18038, p-value = 0.08989
alternative hypothesis: true location shift is greater than 0
```

```
[1] "neuron:  JHNR0003"

   Wilcoxon rank sum test with continuity correction

data:  var by group
W = 14120, p-value = 0.005714
alternative hypothesis: true location shift is greater than 0

[1] "neuron:  JHNR0004"

   Wilcoxon rank sum test with continuity correction

data:  var by group
W = 19875, p-value = 0.1046
alternative hypothesis: true location shift is greater than 0

[1] "neuron:  JINR0002"

   Wilcoxon rank sum test with continuity correction

data:  var by group
W = 14070, p-value = 0.04975
alternative hypothesis: true location shift is less than 0

[1] "neuron:  JINR0012"

   Wilcoxon rank sum test with continuity correction

data:  var by group
W = 2524, p-value = 0.1645
alternative hypothesis: true location shift is greater than 0

[1] "neuron:  JINR0014"

   Wilcoxon rank sum test with continuity correction

data:  var by group
W = 18319, p-value = 0.01627
alternative hypothesis: true location shift is greater than 0

[1] "neuron:  JINR0015"

   Wilcoxon rank sum test with continuity correction

data:  var by group
W = 20908, p-value = 0.0007965
alternative hypothesis: true location shift is greater than 0

[1] "neuron:  JINR0016"

   Wilcoxon rank sum test with continuity correction

data:  var by group
W = 19164, p-value = 0.1981
alternative hypothesis: true location shift is greater than 0

[1] "neuron:  YANR0005"

   Wilcoxon rank sum test with continuity correction

data:  var by group
W = 10413, p-value = 0.324
alternative hypothesis: true location shift is less than 0

[1] "neuron:  YANR0006"
```

```
	Wilcoxon rank sum test with continuity correction

data:  var by group
W = 4139, p-value = 0.0001003
alternative hypothesis: true location shift is less than 0
```

**1.B Effect size of the 27 neurons with alpha < 0.05**

```
 [1] "neuron:  AFNR0003"
# A tibble: 1 × 7
  .y.   group1 group2 effsize    n1    n2 magnitude
* <chr> <chr>  <chr>    <dbl> <int> <int> <ord>    
1 var   LFP    noLFP    0.372   163   157 moderate 
[1] "neuron:  AGNR0002"
# A tibble: 1 × 7
  .y.   group1 group2 effsize    n1    n2 magnitude
* <chr> <chr>  <chr>    <dbl> <int> <int> <ord>    
1 var   LFP    noLFP    0.154    83   237 small    
[1] "neuron:  AGNR0003"
# A tibble: 1 × 7
  .y.   group1 group2 effsize    n1    n2 magnitude
* <chr> <chr>  <chr>    <dbl> <int> <int> <ord>    
1 var   LFP    noLFP    0.190   134   426 small    
[1] "neuron:  AGNR0005"
# A tibble: 1 × 7
  .y.   group1 group2 effsize    n1    n2 magnitude
* <chr> <chr>  <chr>    <dbl> <int> <int> <ord>    
1 var   LFP    noLFP    0.110   177   383 small    
[1] "neuron:  AKNR0000"
# A tibble: 1 × 7
  .y.   group1 group2 effsize    n1    n2 magnitude
* <chr> <chr>  <chr>    <dbl> <int> <int> <ord>    
1 var   LFP    noLFP    0.105   113   447 small    
[1] "neuron:  AKNR0001"
# A tibble: 1 × 7
  .y.   group1 group2 effsize    n1    n2 magnitude
* <chr> <chr>  <chr>    <dbl> <int> <int> <ord>    
1 var   LFP    noLFP   0.0787    98   462 small    
[1] "neuron:  AKNR0004"
# A tibble: 1 × 7
  .y.   group1 group2 effsize    n1    n2 magnitude
* <chr> <chr>  <chr>    <dbl> <int> <int> <ord>    
1 var   LFP    noLFP    0.208    17   123 small    
[1] "neuron:  AKNR0005"
# A tibble: 1 × 7
  .y.   group1 group2 effsize    n1    n2 magnitude
* <chr> <chr>  <chr>    <dbl> <int> <int> <ord>    
1 var   LFP    noLFP   0.0875    73   487 small    
[1] "neuron:  ALNR0003"
# A tibble: 1 × 7
  .y.   group1 group2 effsize    n1    n2 magnitude
* <chr> <chr>  <chr>    <dbl> <int> <int> <ord>    
1 var   LFP    noLFP    0.105   126   436 small    
[1] "neuron:  ALNR0004"
# A tibble: 1 × 7
  .y.   group1 group2 effsize    n1    n2 magnitude
* <chr> <chr>  <chr>    <dbl> <int> <int> <ord>    
1 var   LFP    noLFP    0.189    79   361 small    
[1] "neuron:  ALNR0006"
# A tibble: 1 × 7
  .y.   group1 group2 effsize    n1    n2 magnitude
* <chr> <chr>  <chr>    <dbl> <int> <int> <ord>    
1 var   LFP    noLFP    0.117   163   617 small    
```

```
[1] "neuron:  JANR0002"
# A tibble: 1 × 7
  .y.   group1 group2 effsize    n1    n2 magnitude
* <chr> <chr>  <chr>    <dbl> <int> <int> <ord>    
1 var   LFP    noLFP    0.117    30   190 small    
[1] "neuron:  JANR0006"
# A tibble: 1 × 7
  .y.   group1 group2 effsize    n1    n2 magnitude
* <chr> <chr>  <chr>    <dbl> <int> <int> <ord>    
1 var   LFP    noLFP    0.136   174   216 small    
[1] "neuron:  JANR0007"
# A tibble: 1 × 7
  .y.   group1 group2 effsize    n1    n2 magnitude
* <chr> <chr>  <chr>    <dbl> <int> <int> <ord>    
1 var   LFP    noLFP    0.193   157   233 small    
[1] "neuron:  JANR0008"
# A tibble: 1 × 7
  .y.   group1 group2 effsize    n1    n2 magnitude
* <chr> <chr>  <chr>    <dbl> <int> <int> <ord>    
1 var   LFP    noLFP    0.134   150   240 small    
[1] "neuron:  JANR0009"
# A tibble: 1 × 7
  .y.   group1 group2 effsize    n1    n2 magnitude
* <chr> <chr>  <chr>    <dbl> <int> <int> <ord>    
1 var   LFP    noLFP    0.116   120   270 small    
[1] "neuron:  JANR0011"
# A tibble: 1 × 7
  .y.   group1 group2 effsize    n1    n2 magnitude
* <chr> <chr>  <chr>    <dbl> <int> <int> <ord>    
1 var   LFP    noLFP    0.298   170   220 small    
[1] "neuron:  JANR0012"
# A tibble: 1 × 7
  .y.   group1 group2 effsize    n1    n2 magnitude
* <chr> <chr>  <chr>    <dbl> <int> <int> <ord>    
1 var   LFP    noLFP    0.224   154   236 small    
[1] "neuron:  JANR0017"
# A tibble: 1 × 7
  .y.   group1 group2 effsize    n1    n2 magnitude
* <chr> <chr>  <chr>    <dbl> <int> <int> <ord>    
1 var   LFP    noLFP    0.173   138   252 small    
[1] "neuron:  JDNR0002"
# A tibble: 1 × 7
  .y.   group1 group2 effsize    n1    n2 magnitude
* <chr> <chr>  <chr>    <dbl> <int> <int> <ord>    
1 var   LFP    noLFP    0.128    98   292 small    
[1] "neuron:  JENR0002"
# A tibble: 1 × 7
  .y.   group1 group2 effsize    n1    n2 magnitude
* <chr> <chr>  <chr>    <dbl> <int> <int> <ord>    
1 var   LFP    noLFP    0.235    28    72 small    
[1] "neuron:  JGNR0006"
# A tibble: 1 × 7
  .y.   group1 group2 effsize    n1    n2 magnitude
* <chr> <chr>  <chr>    <dbl> <int> <int> <ord>    
1 var   LFP    noLFP   0.0961   130   260 small    
[1] "neuron:  JGNR0007"
# A tibble: 1 × 7
  .y.   group1 group2 effsize    n1    n2 magnitude
* <chr> <chr>  <chr>    <dbl> <int> <int> <ord>    
1 var   LFP    noLFP    0.121   139   231 small    
[1] "neuron:  JHNR0003"
# A tibble: 1 × 7
  .y.   group1 group2 effsize    n1    n2 magnitude
* <chr> <chr>  <chr>    <dbl> <int> <int> <ord>    
1 var   LFP    noLFP    0.128    79   311 small    
```

```
 [1] "neuron:  JINR0014"
 # A tibble: 1 × 7
   .y.   group1 group2 effsize    n1    n2 magnitude
 * <chr> <chr>  <chr>    <dbl> <int> <int> <ord>
 1 var   LFP    noLFP    0.108   122   268 small
 [1] "neuron:  JINR0015"
 # A tibble: 1 × 7
   .y.   group1 group2 effsize    n1    n2 magnitude
 * <chr> <chr>  <chr>    <dbl> <int> <int> <ord>
 1 var   LFP    noLFP    0.160   153   237 small
 [1] "neuron:  YANR0006"
 # A tibble: 1 × 7
   .y.   group1 group2 effsize    n1    n2 magnitude
 * <chr> <chr>  <chr>    <dbl> <int> <int> <ord>
 1 var   LFP    noLFP    0.257    86   123 small
```

**2.A Mann-Whitney U test results 2/3 - LFP with ECoG-SPW vs. LFP without ECoG-SPW**

```
[1] "neuron:  AFNR0002"

   Wilcoxon rank sum test with continuity correction

data:  var by group
W = 354.5, p-value = 0.08031
alternative hypothesis: true location shift is greater than 0

[1] "neuron:  AFNR0003"

   Wilcoxon rank sum test with continuity correction

data:  var by group
W = 2277.5, p-value = 0.3984
alternative hypothesis: true location shift is greater than 0

[1] "neuron:  AGNR0002"

   Wilcoxon rank sum test with continuity correction

data:  var by group
W = 840.5, p-value = 0.3783
alternative hypothesis: true location shift is less than 0

[1] "neuron:  AGNR0003"

   Wilcoxon rank sum test with continuity correction

data:  var by group
W = 2147, p-value = 0.3353
alternative hypothesis: true location shift is less than 0

[1] "neuron:  AGNR0004"

   Wilcoxon rank sum test with continuity correction

data:  var by group
W = 1826, p-value = 0.04748
alternative hypothesis: true location shift is less than 0

[1] "neuron:  AGNR0005"

   Wilcoxon rank sum test with continuity correction

data:  var by group
W = 2740, p-value = 0.01224
```

```
alternative hypothesis: true location shift is less than 0

[1] "neuron:  AKNR0000"

   Wilcoxon rank sum test with continuity correction

data:  var by group
W = 1084, p-value = 0.574
alternative hypothesis: true location shift is less than 0

[1] "neuron:  AKNR0001"

   Wilcoxon rank sum test with continuity correction

data:  var by group
W = 1147, p-value = 0.07322
alternative hypothesis: true location shift is greater than 0

[1] "neuron:  AKNR0005"

   Wilcoxon rank sum test with continuity correction

data:  var by group
W = 546, p-value = 0.1126
alternative hypothesis: true location shift is less than 0

[1] "neuron:  ALNR0003"

   Wilcoxon rank sum test with continuity correction

data:  var by group
W = 1917, p-value = 0.3317
alternative hypothesis: true location shift is less than 0

[1] "neuron:  ALNR0006"

   Wilcoxon rank sum test with continuity correction

data:  var by group
W = 3362, p-value = 0.4051
alternative hypothesis: true location shift is greater than 0

[1] "neuron:  ANNR0005"

   Wilcoxon rank sum test with continuity correction

data:  var by group
W = 402, p-value = 0.2081
alternative hypothesis: true location shift is greater than 0

[1] "neuron:  JANR0002"

   Wilcoxon rank sum test with continuity correction

data:  var by group
W = 95, p-value = 0.5
alternative hypothesis: true location shift is greater than 0

[1] "neuron:  JANR0004"

   Wilcoxon rank sum test with continuity correction

data:  var by group
W = 5096, p-value = 0.00171
alternative hypothesis: true location shift is greater than 0
```

```
[1] "neuron:  JANR0005"

   Wilcoxon rank sum test with continuity correction

data:  var by group
W = 393.5, p-value = 0.5868
alternative hypothesis: true location shift is greater than 0

[1] "neuron:  JANR0006"

   Wilcoxon rank sum test with continuity correction

data:  var by group
W = 3281.5, p-value = 0.1624
alternative hypothesis: true location shift is less than 0

[1] "neuron:  JANR0007"

   Wilcoxon rank sum test with continuity correction

data:  var by group
W = 2650, p-value = 0.1532
alternative hypothesis: true location shift is less than 0

[1] "neuron:  JANR0008"

   Wilcoxon rank sum test with continuity correction

data:  var by group
W = 2917.5, p-value = 0.6849
alternative hypothesis: true location shift is less than 0

[1] "neuron:  JANR0009"

   Wilcoxon rank sum test with continuity correction

data:  var by group
W = 1971, p-value = 0.08366
alternative hypothesis: true location shift is greater than 0

[1] "neuron:  JANR0010"

   Wilcoxon rank sum test with continuity correction

data:  var by group
W = 2864, p-value = 0.2555
alternative hypothesis: true location shift is less than 0

[1] "neuron:  JANR0011"

   Wilcoxon rank sum test with continuity correction

data:  var by group
W = 3542.5, p-value = 0.4536
alternative hypothesis: true location shift is less than 0

[1] "neuron:  JANR0012"

   Wilcoxon rank sum test with continuity correction

data:  var by group
W = 3069.5, p-value = 0.05283
alternative hypothesis: true location shift is greater than 0

[1] "neuron:  JANR0017"
```

```
   Wilcoxon rank sum test with continuity correction

data:  var by group
W = 2319.5, p-value = 0.5218
alternative hypothesis: true location shift is less than 0

[1] "neuron:  JBNR0005"

   Wilcoxon rank sum test with continuity correction

data:  var by group
W = 397, p-value = 0.1538
alternative hypothesis: true location shift is greater than 0

[1] "neuron:  JENR0003"

   Wilcoxon rank sum test with continuity correction

data:  var by group
W = 887.5, p-value = 0.1454
alternative hypothesis: true location shift is greater than 0

[1] "neuron:  JENR0004"

   Wilcoxon rank sum test with continuity correction

data:  var by group
W = 420.5, p-value = 0.3687
alternative hypothesis: true location shift is greater than 0

[1] "neuron:  JGNR0006"

   Wilcoxon rank sum test with continuity correction

data:  var by group
W = 1396, p-value = 0.0005561
alternative hypothesis: true location shift is greater than 0

[1] "neuron:  JGNR0007"

   Wilcoxon rank sum test with continuity correction

data:  var by group
W = 1142, p-value = 0.05986
alternative hypothesis: true location shift is greater than 0

[1] "neuron:  JGNR0008"

   Wilcoxon rank sum test with continuity correction

data:  var by group
W = 1212.5, p-value = 0.0281
alternative hypothesis: true location shift is greater than 0

[1] "neuron:  JGNR0009"

   Wilcoxon rank sum test with continuity correction

data:  var by group
W = 1864.5, p-value = 0.008071
alternative hypothesis: true location shift is greater than 0

[1] "neuron:  JHNR0003"
```

```
   Wilcoxon rank sum test with continuity correction

data:  var by group
W = 500, p-value = 0.09108
alternative hypothesis: true location shift is less than 0

[1] "neuron:  JHNR0004"

   Wilcoxon rank sum test with continuity correction

data:  var by group
W = 1848, p-value = 0.04604
alternative hypothesis: true location shift is less than 0

[1] "neuron:  JINR0012"

   Wilcoxon rank sum test with continuity correction

data:  var by group
W = 180, p-value = 0.5696
alternative hypothesis: true location shift is greater than 0

[1] "neuron:  JINR0014"

   Wilcoxon rank sum test with continuity correction

data:  var by group
W = 2213.5, p-value = 0.0173
alternative hypothesis: true location shift is greater than 0

[1] "neuron:  JINR0015"

   Wilcoxon rank sum test with continuity correction

data:  var by group
W = 3177, p-value = 0.1452
alternative hypothesis: true location shift is greater than 0

[1] "neuron:  JINR0016"

   Wilcoxon rank sum test with continuity correction

data:  var by group
W = 3362, p-value = 0.3136
alternative hypothesis: true location shift is greater than 0

[1] "neuron:  YANR0005"

   Wilcoxon rank sum test with continuity correction

data:  var by group
W = 1321, p-value = 0.1006
alternative hypothesis: true location shift is less than 0

[1] "neuron:  YANR0006"

   Wilcoxon rank sum test with continuity correction

data:  var by group
W = 799, p-value = 0.04018
alternative hypothesis: true location shift is less than 0
```

**2.B Effect size of the 9 neurons with alpha < 0.05**

```
 [1] "neuron:  AGNR0004"
# A tibble: 1 × 7
  .y.   group1 group2 effsize    n1    n2 magnitude
* <chr> <chr>  <chr>    <dbl> <int> <int> <ord>    
1 var   ECoG   noECoG   0.141    47    93 small    
[1] "neuron:  AGNR0005"
# A tibble: 1 × 7
  .y.   group1 group2 effsize    n1    n2 magnitude
* <chr> <chr>  <chr>    <dbl> <int> <int> <ord>    
1 var   ECoG   noECoG   0.169    55   122 small    
[1] "neuron:  JANR0004"
# A tibble: 1 × 7
  .y.   group1 group2 effsize    n1    n2 magnitude
* <chr> <chr>  <chr>    <dbl> <int> <int> <ord>    
1 var   ECoG   noECoG   0.212    73   117 small    
[1] "neuron:  JGNR0006"
# A tibble: 1 × 7
  .y.   group1 group2 effsize    n1    n2 magnitude
* <chr> <chr>  <chr>    <dbl> <int> <int> <ord>    
1 var   ECoG   noECoG   0.286    17   113 small    
[1] "neuron:  JGNR0008"
# A tibble: 1 × 7
  .y.   group1 group2 effsize    n1    n2 magnitude
* <chr> <chr>  <chr>    <dbl> <int> <int> <ord>    
1 var   ECoG   noECoG   0.187    24    81 small    
[1] "neuron:  JGNR0009"
# A tibble: 1 × 7
  .y.   group1 group2 effsize    n1    n2 magnitude
* <chr> <chr>  <chr>    <dbl> <int> <int> <ord>    
1 var   ECoG   noECoG   0.213    30    98 small    
[1] "neuron:  JHNR0004"
# A tibble: 1 × 7
  .y.   group1 group2 effsize    n1    n2 magnitude
* <chr> <chr>  <chr>    <dbl> <int> <int> <ord>    
1 var   ECoG   noECoG   0.131    34   132 small    
[1] "neuron:  JINR0014"
# A tibble: 1 × 7
  .y.   group1 group2 effsize    n1    n2 magnitude
* <chr> <chr>  <chr>    <dbl> <int> <int> <ord>    
1 var   ECoG   noECoG   0.192    68    54 small    
[1] "neuron:  YANR0006"
# A tibble: 1 × 7
  .y.   group1 group2 effsize    n1    n2 magnitude
* <chr> <chr>  <chr>    <dbl> <int> <int> <ord>    
1 var   ECoG   noECoG   0.189    36    50 small    
```

### 3.A Mann-Whitney U test results 3/3 – LFP-SPWs with no spike vs. all LFP-SPWs

```
[1] "neuron:  AFNR0002"

	Wilcoxon rank sum test with continuity correction

data:  var by group
W = 1205, p-value = 0.9145
alternative hypothesis: true location shift is not equal to 0

[1] "neuron:  AFNR0003"

	Wilcoxon rank sum test with continuity correction

data:  var by group
W = 9516, p-value = 0.5421
```

```
alternative hypothesis: true location shift is not equal to 0

[1] "neuron:  AGNR0002"

   Wilcoxon rank sum test with continuity correction

data:  var by group
W = 3009, p-value = 0.6933
alternative hypothesis: true location shift is not equal to 0

[1] "neuron:  AGNR0003"

   Wilcoxon rank sum test with continuity correction

data:  var by group
W = 2857, p-value = 0.003835
alternative hypothesis: true location shift is not equal to 0

[1] "neuron:  AGNR0004"

   Wilcoxon rank sum test with continuity correction

data:  var by group
W = 6665.5, p-value = 0.5306
alternative hypothesis: true location shift is not equal to 0

[1] "neuron:  AGNR0005"

   Wilcoxon rank sum test with continuity correction

data:  var by group
W = 5875, p-value = 0.01283
alternative hypothesis: true location shift is not equal to 0

[1] "neuron:  AKNR0000"

   Wilcoxon rank sum test with continuity correction

data:  var by group
W = 5320, p-value = 0.6341
alternative hypothesis: true location shift is not equal to 0

[1] "neuron:  AKNR0001"

   Wilcoxon rank sum test with continuity correction

data:  var by group
W = 3942, p-value = 0.585
alternative hypothesis: true location shift is not equal to 0

[1] "neuron:  AKNR0004"

   Wilcoxon rank sum test with continuity correction

data:  var by group
W = 105, p-value = 0.8983
alternative hypothesis: true location shift is not equal to 0

[1] "neuron:  AKNR0005"

   Wilcoxon rank sum test with continuity correction

data:  var by group
W = 1641.5, p-value = 0.7986
alternative hypothesis: true location shift is not equal to 0
```

```
[1] "neuron:  ALNR0003"

   Wilcoxon rank sum test with continuity correction

data:  var by group
W = 4601, p-value = 0.8349
alternative hypothesis: true location shift is not equal to 0

[1] "neuron:  ALNR0004"

   Wilcoxon rank sum test with continuity correction

data:  var by group
W = 1672.5, p-value = 0.8546
alternative hypothesis: true location shift is not equal to 0

[1] "neuron:  ALNR0006"

   Wilcoxon rank sum test with continuity correction

data:  var by group
W = 10278, p-value = 0.7602
alternative hypothesis: true location shift is not equal to 0

[1] "neuron:  ANNR0003"

   Wilcoxon rank sum test with continuity correction

data:  var by group
W = 967, p-value = 0.8253
alternative hypothesis: true location shift is not equal to 0

[1] "neuron:  ANNR0005"

   Wilcoxon rank sum test with continuity correction

data:  var by group
W = 1843, p-value = 0.9151
alternative hypothesis: true location shift is not equal to 0

[1] "neuron:  JANR0001"

   Wilcoxon rank sum test with continuity correction

data:  var by group
W = 1217, p-value = 0.8166
alternative hypothesis: true location shift is not equal to 0

[1] "neuron:  JANR0002"

   Wilcoxon rank sum test with continuity correction

data:  var by group
W = 370, p-value = 0.82
alternative hypothesis: true location shift is not equal to 0

[1] "neuron:  JANR0004"

   Wilcoxon rank sum test with continuity correction

data:  var by group
W = 13132, p-value = 0.3338
alternative hypothesis: true location shift is not equal to 0

[1] "neuron:  JANR0005"
```

```
   Wilcoxon rank sum test with continuity correction

data:  var by group
W = 1507, p-value = 0.668
alternative hypothesis: true location shift is not equal to 0

[1] "neuron:  JANR0006"

   Wilcoxon rank sum test with continuity correction

data:  var by group
W = 6353.5, p-value = 0.9967
alternative hypothesis: true location shift is not equal to 0

[1] "neuron:  JANR0007"

   Wilcoxon rank sum test with continuity correction

data:  var by group
W = 5303.5, p-value = 0.2554
alternative hypothesis: true location shift is not equal to 0

[1] "neuron:  JANR0008"

   Wilcoxon rank sum test with continuity correction

data:  var by group
W = 7319, p-value = 0.3271
alternative hypothesis: true location shift is not equal to 0

[1] "neuron:  JANR0009"

   Wilcoxon rank sum test with continuity correction

data:  var by group
W = 3457, p-value = 0.1747
alternative hypothesis: true location shift is not equal to 0

[1] "neuron:  JANR0010"

   Wilcoxon rank sum test with continuity correction

data:  var by group
W = 5207.5, p-value = 0.4903
alternative hypothesis: true location shift is not equal to 0

[1] "neuron:  JANR0011"

   Wilcoxon rank sum test with continuity correction

data:  var by group
W = 9583.5, p-value = 0.07696
alternative hypothesis: true location shift is not equal to 0

[1] "neuron:  JANR0012"

   Wilcoxon rank sum test with continuity correction

data:  var by group
W = 2748.5, p-value = 0.05273
alternative hypothesis: true location shift is not equal to 0

[1] "neuron:  JANR0017"

   Wilcoxon rank sum test with continuity correction
```

```
data:  var by group
W = 4746, p-value = 0.46
alternative hypothesis: true location shift is not equal to 0

[1] "neuron:  JBNR0003"

	Wilcoxon rank sum test with continuity correction

data:  var by group
W = 556.5, p-value = 0.4042
alternative hypothesis: true location shift is not equal to 0

[1] "neuron:  JBNR0005"

	Wilcoxon rank sum test with continuity correction

data:  var by group
W = 1625, p-value = 0.9043
alternative hypothesis: true location shift is not equal to 0

[1] "neuron:  JDNR0001"

	Wilcoxon rank sum test with continuity correction

data:  var by group
W = 4844, p-value = 0.6455
alternative hypothesis: true location shift is not equal to 0

[1] "neuron:  JDNR0002"

	Wilcoxon rank sum test with continuity correction

data:  var by group
W = 4459.5, p-value = 0.4333
alternative hypothesis: true location shift is not equal to 0

[1] "neuron:  JENR0001"

	Wilcoxon rank sum test with continuity correction

data:  var by group
W = 2671.5, p-value = 0.3856
alternative hypothesis: true location shift is not equal to 0

[1] "neuron:  JENR0002"

	Wilcoxon rank sum test with continuity correction

data:  var by group
W = 268.5, p-value = 0.9577
alternative hypothesis: true location shift is not equal to 0

[1] "neuron:  JENR0003"

	Wilcoxon rank sum test with continuity correction

data:  var by group
W = 2210, p-value = 0.7384
alternative hypothesis: true location shift is not equal to 0

[1] "neuron:  JENR0004"

	Wilcoxon rank sum test with continuity correction
```

```
data:  var by group
W = 1023.5, p-value = 0.3151
alternative hypothesis: true location shift is not equal to 0

[1] "neuron:  JGNR0006"

   Wilcoxon rank sum test with continuity correction

data:  var by group
W = 6595, p-value = 0.2082
alternative hypothesis: true location shift is not equal to 0

[1] "neuron:  JGNR0007"

   Wilcoxon rank sum test with continuity correction

data:  var by group
W = 5634.5, p-value = 0.2914
alternative hypothesis: true location shift is not equal to 0

[1] "neuron:  JGNR0008"

   Wilcoxon rank sum test with continuity correction

data:  var by group
W = 2643.5, p-value = 0.7381
alternative hypothesis: true location shift is not equal to 0

[1] "neuron:  JGNR0009"

   Wilcoxon rank sum test with continuity correction

data:  var by group
W = 7809, p-value = 0.5282
alternative hypothesis: true location shift is not equal to 0

[1] "neuron:  JHNR0003"

   Wilcoxon rank sum test with continuity correction

data:  var by group
W = 1309.5, p-value = 0.2016
alternative hypothesis: true location shift is not equal to 0

[1] "neuron:  JHNR0004"

   Wilcoxon rank sum test with continuity correction

data:  var by group
W = 923, p-value = 0.2052
alternative hypothesis: true location shift is not equal to 0

[1] "neuron:  JINR0002"

   Wilcoxon rank sum test with continuity correction

data:  var by group
W = 5333, p-value = 0.4141
alternative hypothesis: true location shift is not equal to 0

[1] "neuron:  JINR0012"

   Wilcoxon rank sum test with continuity correction

data:  var by group
W = 577, p-value = 0.5826
```

```
alternative hypothesis: true location shift is not equal to 0

[1] "neuron:  JINR0014"

   Wilcoxon rank sum test with continuity correction

data:  var by group
W = 4180.5, p-value = 0.05133
alternative hypothesis: true location shift is not equal to 0

[1] "neuron:  JINR0015"

   Wilcoxon rank sum test with continuity correction

data:  var by group
W = 7450, p-value = 0.1028
alternative hypothesis: true location shift is not equal to 0

[1] "neuron:  JINR0016"

   Wilcoxon rank sum test with continuity correction

data:  var by group
W = 7393.5, p-value = 0.08824
alternative hypothesis: true location shift is not equal to 0

[1] "neuron:  YANR0005"

   Wilcoxon rank sum test with continuity correction

data:  var by group
W = 4509, p-value = 0.7348
alternative hypothesis: true location shift is not equal to 0

[1] "neuron:  YANR0006"

   Wilcoxon rank sum test with continuity correction

data:  var by group
W = 2738, p-value = 0.3233
alternative hypothesis: true location shift is not equal to 0
```

**3.B Effect size of the 2 neurons with alpha < 0.05**

```
[1] "neuron:  AGNR0003"
# A tibble: 1 × 7
  .y.   group1 group2     effsize    n1    n2 magnitude
* <chr> <chr>  <chr>        <dbl> <int> <int> <ord>
1 var   allLFP LFPnoSpike   0.223   134    34 small
[1] "neuron:  AGNR0005"
# A tibble: 1 × 7
  .y.   group1 group2     effsize    n1    n2 magnitude
* <chr> <chr>  <chr>        <dbl> <int> <int> <ord>
1 var   allLFP LFPnoSpike   0.163   177    56 small
```
